# Phosphodiesterase Type 4 anchoring regulates cAMP signaling to Popeye domain-containing proteins

**DOI:** 10.1101/2020.09.10.290825

**Authors:** Amy J Tibbo, Sara Dobi, Aisling McFall, Gonzalo S Tejeda, Connor Blair, Ruth MacLeod, Niall MacQuaide, Caglar Gök, William Fuller, Brian O Smith, Godfrey L Smith, Thomas Brand, George S Baillie

## Abstract

Cyclic AMP is a ubiquitous second messenger used to transduce intracellular signals from a variety of Gs-coupled receptors. Compartmentalisation of protein intermediates within the cAMP signaling pathway underpins receptor-specific responses. The cAMP effector proteins protein-kinase A and EPAC are found in complexes that also contain phosphodiesterases whose presence ensures a coordinated cellular response to receptor activation events. Popeye proteins are the most recent class of cAMP effectors to be identified and have crucial roles in cardiac pacemaking and conduction. We report the first observation that Popeye proteins exist in complexes with members of the PDE4 family in cardiac myocytes thus restricting cAMP signaling. We show that POPDC1 preferentially binds the PDE4A sub-family via a specificity motif in the PDE4 UCR1 region and that PDE4s bind to the Popeye domain of POPDC1 in a region known to be susceptible to a mutation that causes human disease. Using a cell-permeable disruptor peptide that displaces the POPDC1-PDE4 complex we show that PDE4 activity localized to POPDC1 is essential to maintain action potential duration in beating cardiac myocytes.

## Introduction

Phosphodiesterases (PDEs) are the only known family of enzymes that can desensitize cAMP signaling via their ability to degrade cAMP. As there are 11 different multi-gene families of PDEs that give rise to over 100 isoforms, which are distributed in a tissue- and cell type specific manner, it has been hypothesized that the precise cellular location of each isoform underpins the function of each PDE species (Baillie *et al*, 2019; Maurice *et al*, 2014). A good example that illustrates the concept is the PDE type 4 isoform called PDE4D5 (Wills *et al*, 2016). It can be translocated to the vicinity of G_s_-coupled receptors by the signaling scaffold protein beta-arrestin in order to hydrolyze cAMP, while beta-arrestin concomitantly hinders G-protein signaling (Baillie *et al*, 2003). This dual functionality is essential for effective desensitization of β -adrenoceptor signaling (Willoughby *et al*, 2007). PDE4D5 can also anchor to another scaffold protein, RACK1, to localize the enzyme at the leading edge of moving cells to allow direction sensing (Serrels *et al*, 2010). Specific disruption of the RACK-PDE4D5 complex hinders cell movement via an exchange factor directly activated by cAMP (EPAC)-mediated signaling mechanism but does not affect β -adrenergic signaling (Serrels *et al*, 2011). Conversely, disassembly of the β-arrestin-PDE4D5 complex allows prolonged cAMP signaling following β-adrenergic stimulation that results in hyper-phosphorylation of the β -adrenergic receptor by protein kinase A (PKA) but this action does not affect PDE4D5 located in focal adhesions (Smith *et al*, 2007).

The close relationship between PDE anchoring and appropriate activation of cAMP effectors such as EPAC and PKA has been widely investigated in the last decade (Baillie *et al.*, 2019; Maurice *et al.*, 2014). This is particularly germane in cAMP signaling networks that augment excitation-contraction coupling in the heart during times of increased demand. Highly localized increases in cAMP (Stangherlin *et al*, 2014) that appear in cardiac myocytes following β -adrenergic stimulation activate discrete pools of PKA that phosphorylate calcium handling proteins such as the L-type calcium channel, ryanodine receptor (RYR2) and the SERCA regulator phospholamban (Mika *et al*, 2013). Each of these proteins forms a functional complex with an A-kinase anchoring protein (AKAP, which scaffolds PKA) and a PDE isoform (reviewed in (Maurice *et al.*, 2014)). This arrangement allows a pulse of cAMP to briefly breach the threshold of activation of the anchored PKA within each complex and deliver a transient augmentation of calcium flux following phosphorylation of the respective PKA substrates. The anchored PDE influences the size and duration of the local PKA drive whilst also guarding against inappropriate activation at resting heart rate (Fischmeister *et al*, 2006). There are many other non-cardiac examples of cAMP effectors (PKA, EPAC and cyclic nucleotide-gated (CNG) channels) that exist in complex with PDEs (reviewed in (Maurice *et al.*, 2014)).

An important, but as yet only partially characterized family of plasma membrane-localized cAMP effector proteins was discovered and named the Popeye domain containing (POPDC) proteins due to their abundant expression in the heart and skeletal muscle ((Andree *et al*, 2000);(Froese *et al*, 2012);(Brand & Schindler, 2017). POPDC proteins are mainly localized at the sarcolemma and carrying an evolutionary conserved intracellular Popeye domain, which acts as a high-affinity cAMP binding domain ((Brand, 2018; Froese *et al.*, 2012). Loss-of-function mutations of Popdc1 and Popdc2 in mice and zebrafish established an important role for these genes in cardiac pacemaking and conduction ((Froese *et al.*, 2012; Kirchmaier *et al*, 2012; Schindler *et al*, 2016). Mutations in POPDC1, POPDC2 and POPDC3 have been recently identified in patients suffering from cardiac arrhythmia and/or limb-girdle muscular dystrophy, respectively ((De Ridder *et al*, 2019; Indrawati *et al*, 2020; Schindler *et al.*, 2016; Vissing *et al*, 2019). Moreover, POPDC proteins have also been implicated in atrial fibrillation, long QT syndrome and heart failure (Gingold-Belfer *et al*, 2011; Tan *et al*, 2013; Wang *et al*, 2016). Work in colon cancer and other tumors suggests that POPDC1 may also act as a tumor suppressor (Amunjela *et al*, 2019; Parang *et al*, 2018). Several protein-protein interaction partners of POPDC proteins have been identified including the tandem pore-domain background potassium channel TWIK-related K+ channel 1 (TREK-1). When TREK-1 is co-expressed with POPDC proteins in Xenopus oocytes, the channel displays enhanced membrane trafficking and conductivity which is sensitive to changes in cAMP levels (Froese *et al.*, 2012; Rinne *et al*, 2020; Schindler *et al.*, 2016).

Effector proteins of the cAMP pathway form complexes with PDEs, which limits their activation in response to elevations in cAMP levels. We show here for the first time that POPDC proteins are similar to other cAMP effectors in that they form a complex with PDE4s and, in particular, with members of the PDE4A sub-family. We go on to show that PDE4 activity regulates the ability of POPDC proteins to bind cAMP and to interact with TREK-1. This complex may therefore represent a novel therapeutic target for cardiac arrhythmia, muscular dystrophy and cancer.

## Results

### PDE4 enzymes and POPDC1 co-localise in ventricular myocytes

As PDEs, and in particular PDE4, function as part of “signalosomes” where they restrict the availability of cAMP to cAMP effectors (Fertig & Baillie, 2018), we were interested to see whether the least characterized and newly discovered cAMP effector protein POPDC1 (Brand & Schindler, 2017) also existed in a complex with PDE4. Initial studies in HEK293 cells that were transiently transfected with POPDC1 and PDE4A suggested that the proteins co-localized (Fig. 1A,B) and this prompted us determine whether this was also the case in neonatal rat ventricular myocytes (NRVMs) where POPDC proteins mainly localize to the plasma membrane (Brand, 2018). In agreement with previous work, POPDC1 is concentrated in the plasma membrane of NRVMs (Fig. 1A,C) where it co-localized with a membrane-bound pool of PDE4A. The PDE4A5 (murine orthologue of PDE4A4) enzyme is known to have both membrane and cytosolic pools conferred by different targeting cassettes within its sequence (Beard *et al*, 2002), in agreement with observed staining pattern in NRVMs (Fig. 1A). To verify the specific co-localization of POPDC1 and PDE4A5 we performed in situ proximity ligation assays (PLA), which confirmed the intimate spatial co-segregation of these proteins in HEK293 cells, NRVMs and adult rabbit ventricular myocytes (ARVMs) (Fig. 1D). The segregation of PDE4A and POPDC at the membrane could also be observed in cellular fractionation experiments using transfected HEK293 cells and NRVMs where, as expected, POPDC1 was found to be exclusively membrane bound whereas PDE4A was split between fractions (Figs. 1E and 1F). The integrity of the membrane fraction was validated by using the Na^+^/K^+^-ATPase as a marker.

**Figure 1.**
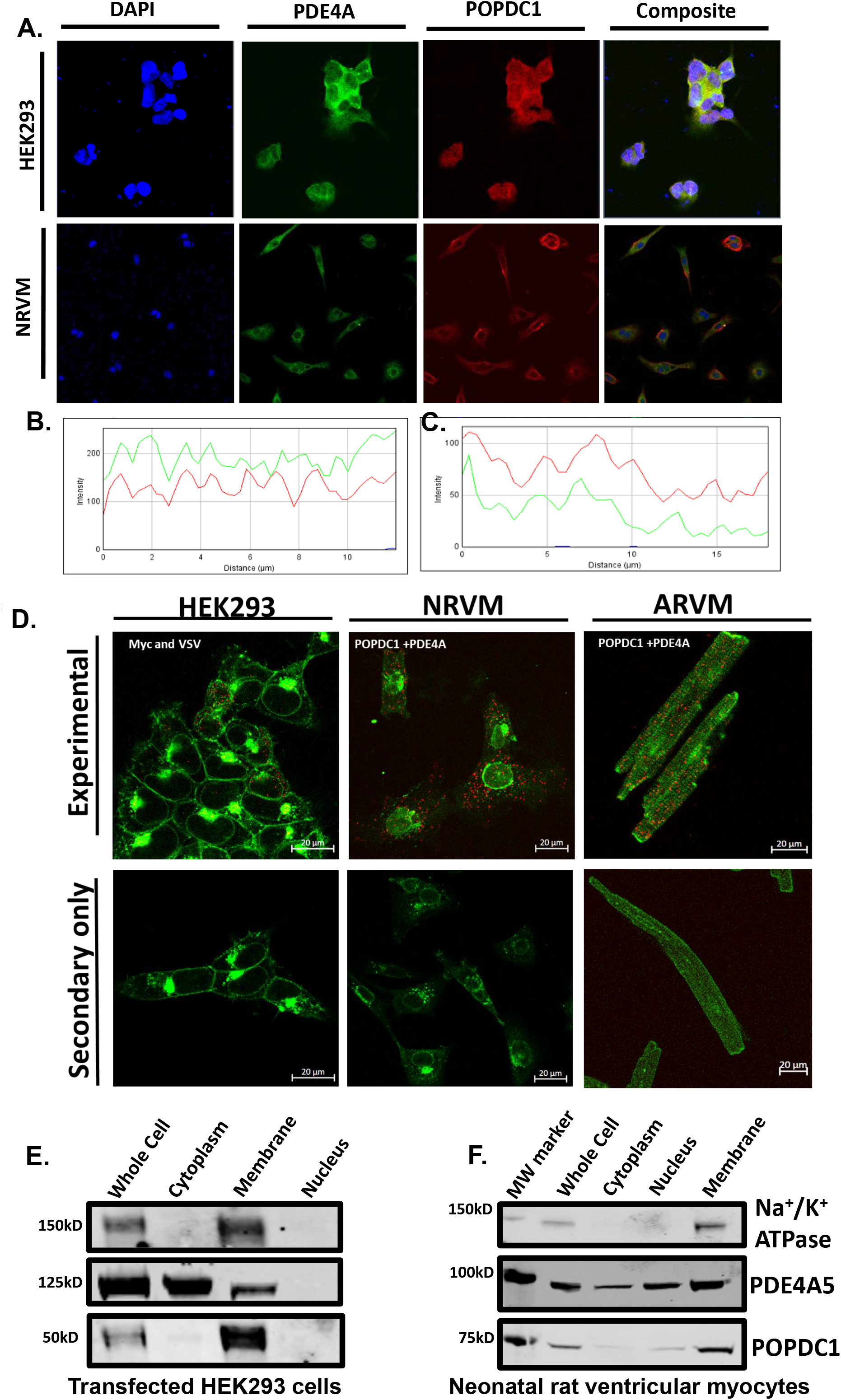
POPDC1 co-localises with PDE4. A. Immunofluorescence showing colocalization of PDE4A4 (green) and POPDC1 (red) in transiently transfected HEK293 and NRVMs. B,C. The panels represent typical line scan intensity profile of (B) transfected HEK293 cells (Pearson’s coefficient = 0.6.) and (C) endogenously expressing NRVMs (Pearson’s coefficient = 0.6). D. Proximity ligation analysis shows that POPDC1 and PDE4 colocalized in transfected HEK293 cells, NRVMs and ARVMs (upper panels). No background was detected in controls where only the secondary antibodies were used (lower panels). E,F. POPDC1 and PDE4A4/5 are detected in the membrane fraction after subcellular fractionation of cell lysate isolated from (E) transfected HEK293 cells or (F) NRVMs.

### POPDC1 interacts directly with PDE4

To determine whether POPDC1 exists in a complex with PDE4 in a similar fashion to other cAMP-effector proteins, we immunoprecipitated transiently transfected MYC-tagged POPDC1 from HEK293 cells and probed for co-transfected VSV-tagged PDE4A5. We successfully pulled down VSV-PDE4A5 with POPDC1 (Fig. 2A) suggesting a close interaction of the exogenous proteins. Reassuringly, we could also recreate these findings in NRVMs expressing both proteins endogenously (Fig. 2B) suggesting that a pre-formed complex between POPDC1 and PDE4 exists in the cardiac myocytes. Next, we sought to determine whether the interaction between the proteins was direct or mediated by a scaffolding protein. We purified MBP-PDE4A4 (human orthologue of PDE4A5) and GST-POPDC1 and incubated them together. Isolation of POPDC1 using the GST tag also co-immunoprecipitated MBP-PDE4A (Fig. 2C). Likewise, purifying PDE4A4 using the MBP-tag, also pulled down GST-POPDC1, suggesting that there is a direct interaction (Fig. 2D). The assumption that the proteins interacted directly was further verified using far-western techniques, in which immobilized GST-coupled versions of known PDE4A binding proteins UBC9 and P75 NTR (Houslay *et al*, 2017) and POPDC1 were overlaid with MBP-PDE4A4 (Fig. 2E). All three proteins bound directly to PDE4A4 (Fig. 2F) whereas a control protein RHE PfPdfx did not (Fig. 2G,H), confirming that POPDC1 is able to bind directly to PDE4A4.

**Figure 2.**
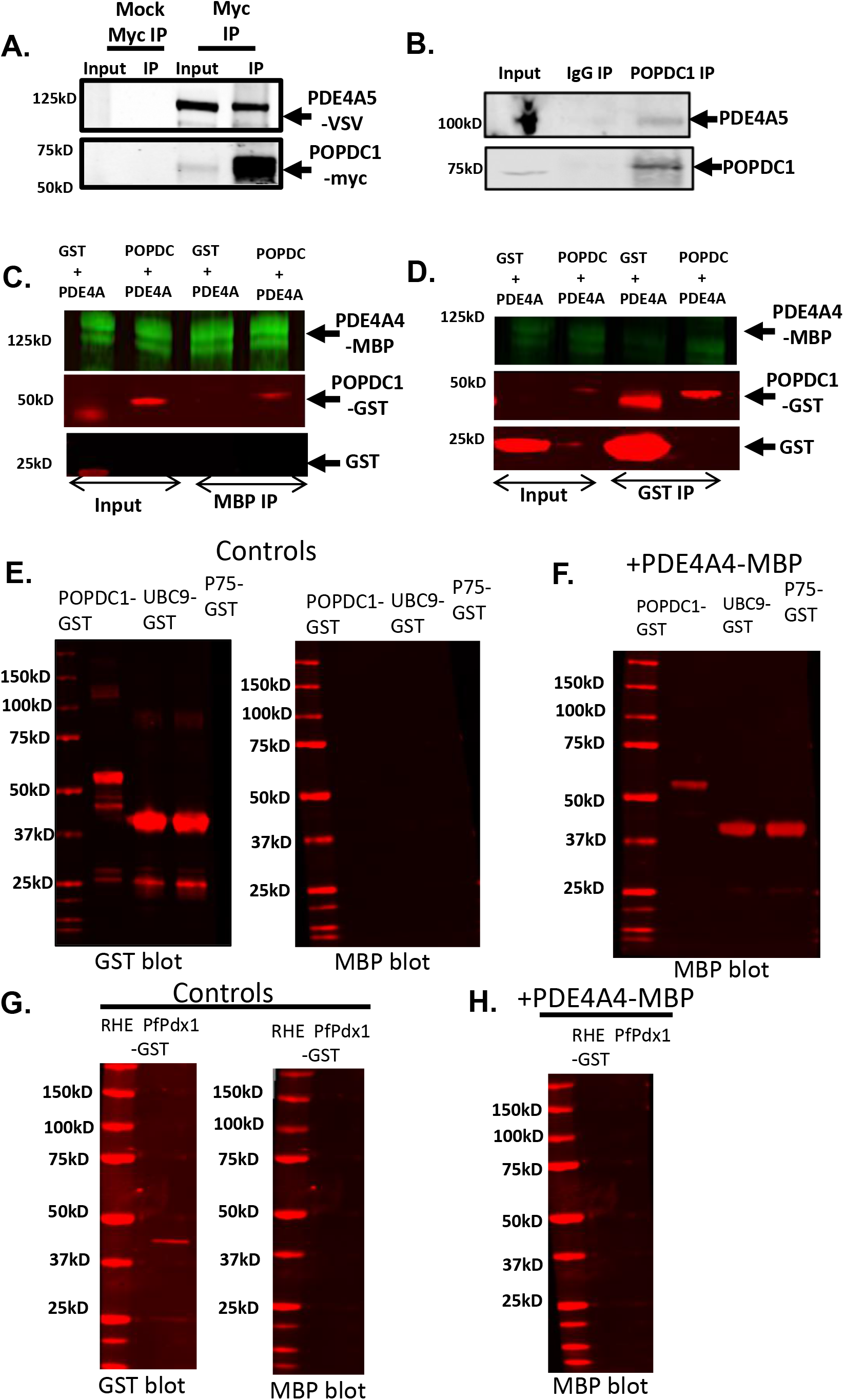
PDE4A and POPDC1 bind directly to each other. A. Co-immunoprecipitation identifying POPDC1 in a complex with PDE4A5 from transiently transfected HEK293 overexpressing PDE4A5-VSV and POPDC1-myc and B. endogenously expressing NRVMs. C. Recombinant purified POPDC1-GST and PDE4A4-MBP were mixed at equal concentrations prior to co-immunoprecipitations using glutathione-agarose beads or D. amylose resin, that binds to the MBP tag. E. Far Western blotting where recombinant purified POPDC1-GST, UBC9-GST and P75-GST proteins were run on SDS-PAGE and blotted for GST, to ensure the proteins had been successfully transferred, and for MBP, to ensure there was no non-specific binding. F. Membranes were then overlaid with recombinant purified PDE4A4-MBP and re-probed with MBP to detect any interaction between the ‘bait’ proteins and the overlaid PDE. G and H. A negative control experiment was carried out in the same manner but utilizing recombinant purified RHE PfPdx1-GST protein as it has not been shown to interact with PDE4A4.

### Mapping the PDE4-POPDC1 interface using peptide array

PDEs are often localized to specific subcellular locations by virtue of protein-protein interactions and peptide array technology has been utilized to define the binding sites that anchor these enzymes (Baillie, 2015; Bolger *et al*, 2006). We used this technique to map the PDE4A binding site on POPDC1. Initial screening of a library of immobilized peptides of human POPDC1 (25mers shifted by 5 amino acids and encompassing the entire human POPDC1 sequence (UniProt sequence: Q8NE79) suggested a PDE4A4 (but not PDE4D9) binding site, which is localized within the Popeye domain (Fig. 3A) encompassing amino acids 154 to 178. Alanine scanning analysis of this 25mer (Fig. 3B) identified 5 essential amino acids in POPDC1 that are required for interaction with PDE4A4, namely R(172), L(173), S(174), L (176) and K (178). The identification of these residues was verified by a motif scanning experiment where the RLSILLK motif (Fig. 3C, in red) was sequentially shifted through arrays of the POPDC1 sequence of interest. All peptides containing an intact RLSILLK motif showed some degree of binding to PDE4A4 (Fig. 3C). N- and C-terminal truncations of the 154-178 POPDC1 25mer sequence also demonstrated the importance of the RLSILLK motif as binding occurred with peptides that were N-terminally truncated but still retained the RLSILLK domain (Fig. 3D), whereas binding was ablated when the C-terminal residue K (179) was deleted (Fig. 3E)

**Figure 1.**
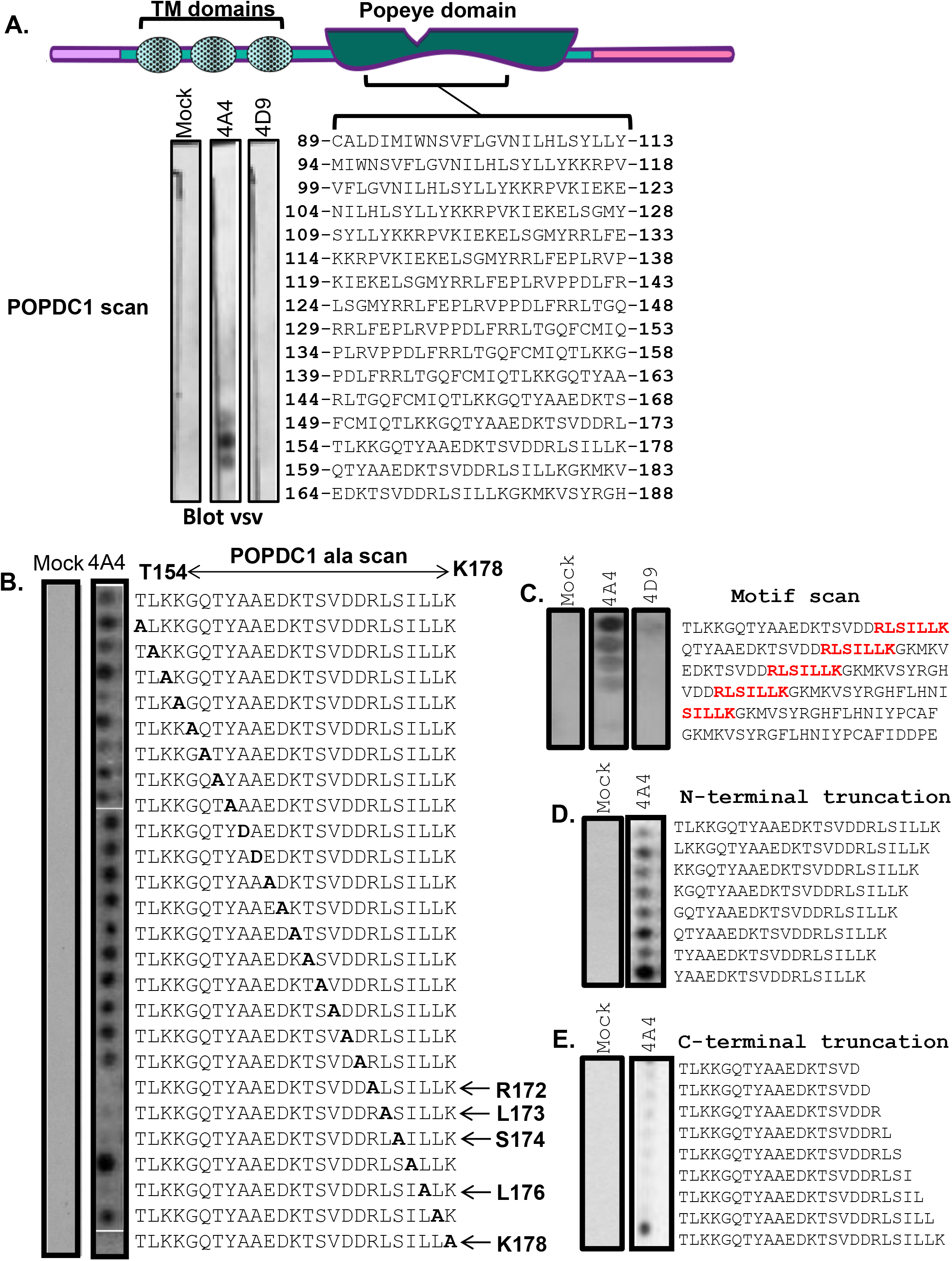
Mapping the PDE4 docking region on POPDC1. A. Peptide arrays encompassing the full sequence of the POPDC1 protein were overlaid with lysate from HEK293 cells over-expressing PDE4A4-VSV or PDE4D9-VSV. Each spot contains an immobilized 25mer sequence derived from the POPDC1 sequence. Each peptide is related to its neighbours by a 5 amino acid shift. Dark spots indicate peptides that interacted with PDE4. B. The initial binding site identified in A. (POPDC1 aa154 – aa178) was further interrogated by alanine scanning to identify single amino acids required for the interaction. C. A motif scan array was constructed to evaluate the importance of the RLSILLK motif discovered in B. This was overlaid with PDE4A4-VSV. D. To confirm whether the binding motif RLSILLK was important for the PDE4 interaction, an N-terminal truncation or E. a C-terminal truncation peptide array was analyzed. For all peptide array experiments negative control experiment was performed using mock transfected HEK293 cell lysate.

PDE4 enzymes are transcribed from 4 genes (A, B, C, and D) that have multiple gene products and in total approximately 25 unique isoforms exist. PDE4 isoforms have non-redundant functions that are related to differential targeting mechanisms, which place them in specific cellular locations(Baillie *et al.*, 2019). As PDE4D9 did not bind to POPDC1 in peptide array experiments (Fig. 3A), we speculated as to whether this represented a PDE4A-specific interaction. Indeed, we were unable to co-immunoprecipitate PDE4B or PDE4D long isoforms with myc-tagged POPDC1 from transfected HEK293 cells (Suppl. Fig. 1A) and PLA experiments in NRVMs showed a significantly higher degree of co-localization between POPDC1 and pan-PDE4A compared with pan-PDE4B and pan-PDE4D antibodies (Suppl. Fig. 1B) even though all subfamilies are expressed to similar levels in this cell type (Johnson *et al*, 2012; Richter *et al*, 2011). On overlaying PDE4AA peptide arrays with purified POPDC1 Popeye domain, we were intrigued to find a POPDC binding domain within the UCR1 region (Fig. 4A). The strongest interaction was detected with a peptide spanning D155 to P180 of PDE4A4 (Fig. 4A), which contains a region whose amino acid sequence is less stringently conserved than most of UCR1 (Fig. 4B). This region has 6 regions of variance which we named “motifs 1-6” (Fig. 4B). A similar experiment overlaying purified full-length POPDC revealed a similar binding domain in UCR1 (Fig. 4E). Alanine scanning of the D155 to P180 25mer revealed an essential region that encompassed K159 to H172 of PDE4A4 (Fig. 4C). This region was also shown to be essential for POPDC1 binding in C-terminal truncation analysis (Fig. 4D). Comparison of the cognate regions taken from the two most highly expressed cardiac PDE4s, PDE4A and PDE4D (Richter *et al.*, 2011) showed that sequences from the PDE4A subfamily gave the most robust interactions with the purified full-length POPDC1 (Fig. 4F, upper two array spots) suggesting that the divergence in the UCR1 sequence in this area underpins POPDC1’s preference for PDE4A over other subfamilies. Sequential substitution of each divergent motif on PDE4A and PDE4D UCR1 sequences showed that POPDC1 binding to PDE4D could be attenuated when motifs 2,3,4 and 5 were mutated but reductions of POPDC1 binding to PDE4A sequences was only observed when motif 3 was substituted (Fig. 4F).

**Figure 2.**
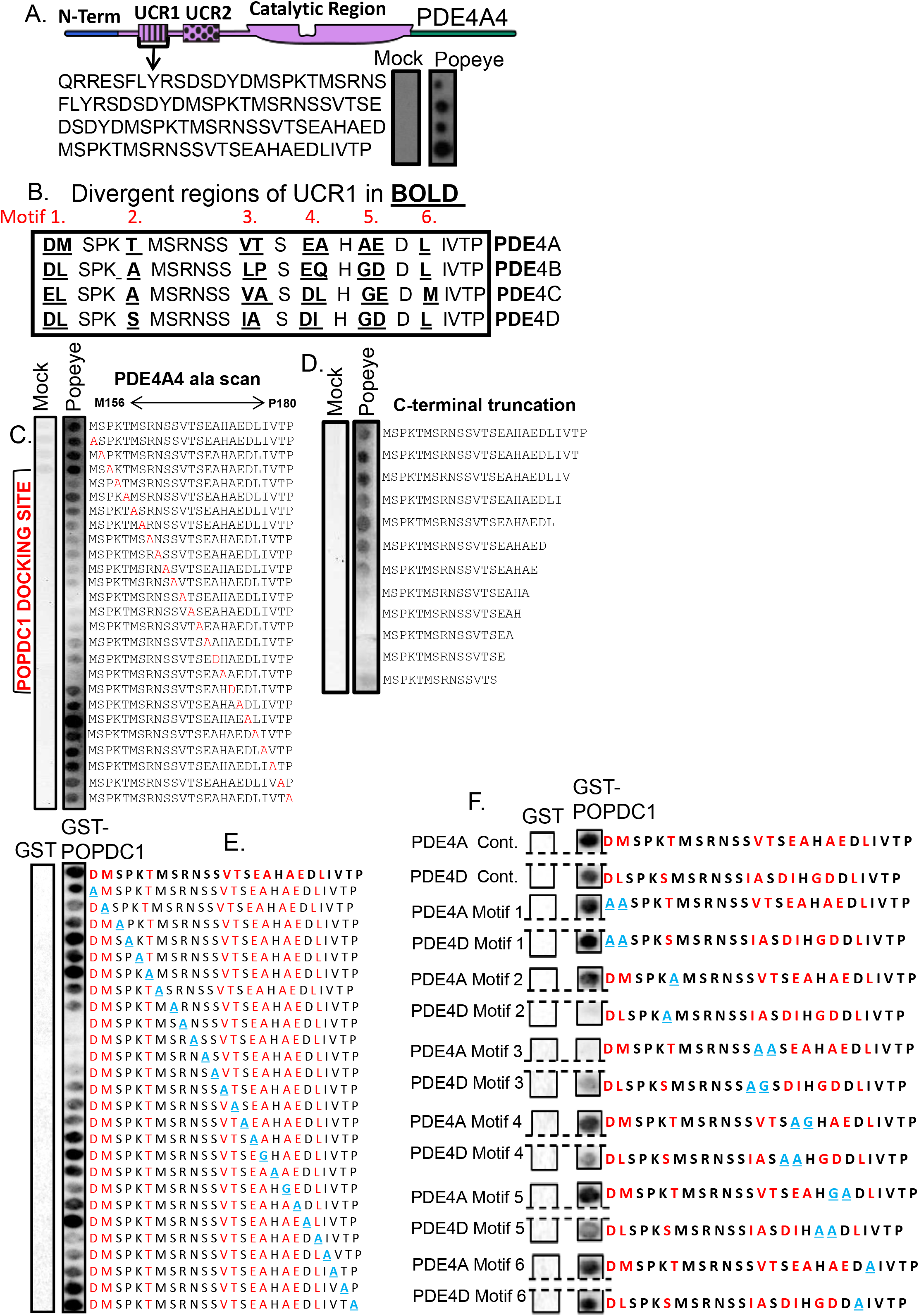
POPDC1 binds to a site within the PDE4 UCR1 domain. A. A peptide array of 25mers shifted by 5 amino acids covering the full PDE4A4 sequence was overlaid with purified recombinant protein consisting of the popeye domain of POPDC1 tagged with GST. A binding region was identified in the UCR1 domain of PDE4A and the modular structure of PDE4A is shown. Mock control was treated under same conditions but with BSA added instead of popeye domain. B. The popeye binding domain in UCR1 contains 6 divergent regions (termed motifs 1-6 in red) when sequences from PDE4 A,B,C and are compared. Divergent amino acids are identified in bold and underlined. C. An alanine scanning array of the popeye binding domain was overlaid with purified recombinant protein consisting of the popeye domain of POPDC1 tagged with GST. Amino acids that are alanised (or changed to Asp if alanine) are shown in red. Control array (Mock) was treated in same manner but the purified popeye domain was replaced with BSA. D. A C-terminal truncation array was overlaid with purified recombinant protein consisting of the popeye domain of POPDC1 tagged with GST. Control array (Mock) was treated in same manner but the purified popeye domain was replaced with BSA. E. An alanine scanning array of the popeye binding domain was overlaid with purified recombinant protein consisting of full length POPDC1 tagged with GST to identify essential amino acids where dark spots are attenuated. Amino acids that are alanised (or changed to glycine if alanine) are shown in light blue. Divergent amino acids are shown in red. Control array (GST) was treated in same manner but the purified popeye domain was replaced with GST. F. Spot arrays that compared POPDC1 binding sites from PDE4A and PDE4D overlaid with purified recombinant protein consisting of full length POPDC1 tagged with GST to identify role of divergent sequences (motifs 1-6). Each of the divergent motif was sequentially alanised (or changed to Glycine if alanine, shown in light blue) and binding of POPDC1 determined. Divergent amino acids are shown in red. Control array (GST) was treated in same manner but the purified popeye domain was replaced with GST

Since UCR1 present in all long PDE4 isoforms, we decided to further investigate the specificity for PDE4A. Data from co-immunoprecipitations performed using transfected HEK293 cells complemented the data from initial peptide array studies (Fig. 3A) that suggested a degree of PDE4A selectivity as neither of the long isoforms PDE4D7 or PDE4B1 was found to interact with POPDC1 (Fig. 5A). As transfected constructs can often mislocalize when over-expressed, the exclusivity of PDE4A binding to POPDC1 was determined using NRVMs and proximity ligation microscopy, a method we have successfully used in the past to confirm co-localisation of endogenous proteins with their binding partners (Sin *et al*, 2015). A robust PLA signal was detected between POPDC1 and PDE4A (Fig. 1D). In comparison to PDE4A, significantly weaker signals were recorded for POPDC1 interactions with endogenous PDE4B and PDE4D were observed (Fig. 5B and 5C) reinforcing the notion that PDE4A was the “preferred” partner due to the optimal POPDC1 binding region in the UCR1 divergent region motif 3 (Fig. 4B). This situation has similarities to the association of PDE4D5 with beta-arrestin-2 where all PDE4s can interact with the scaffold but PDE4D5 is “preferred” due to an optimized binding motif within the PDE’s N-terminal region (Wills *et al.*, 2016).

**Figure 3.**
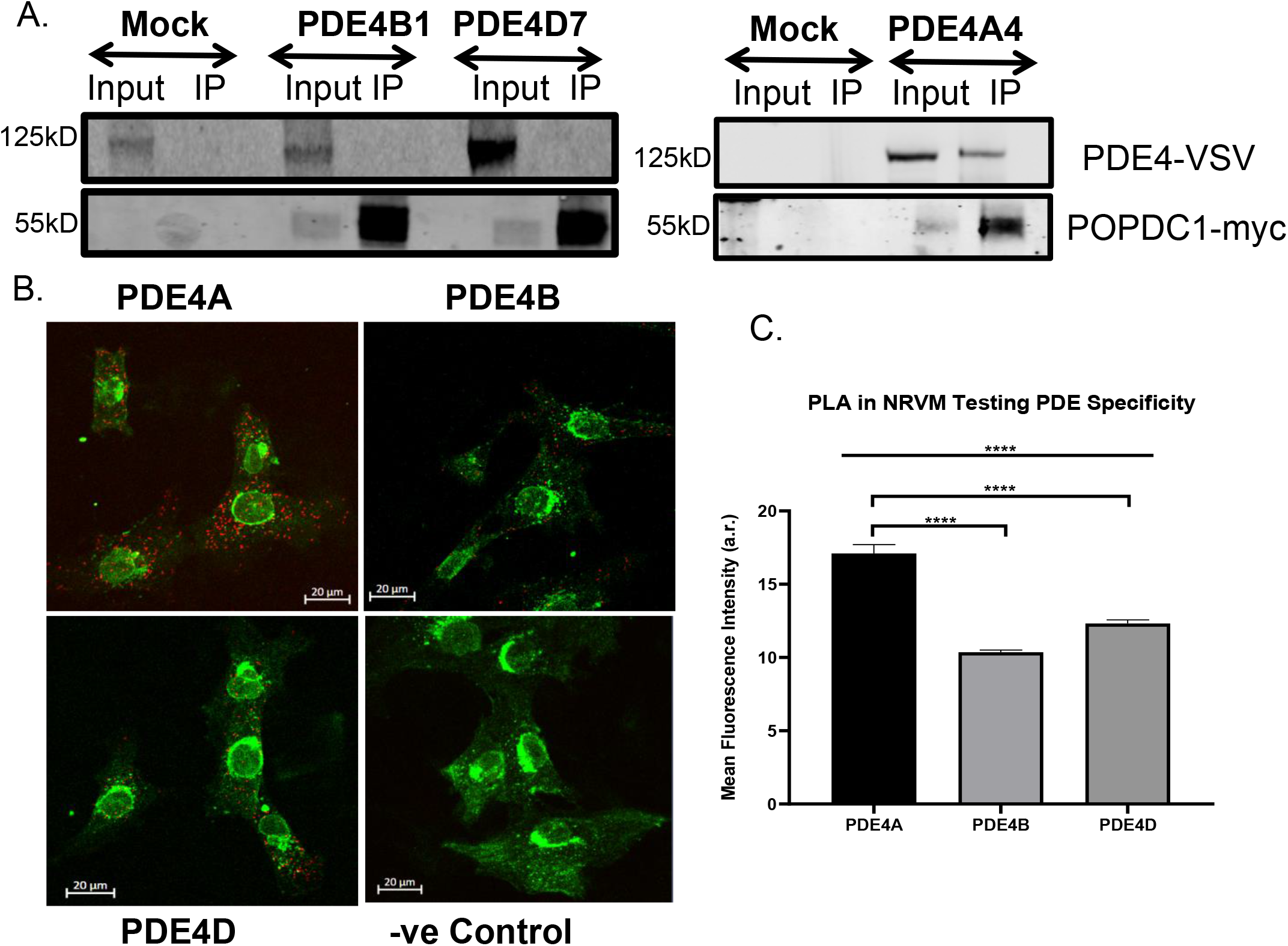
POPDC1 interacts preferentially with the PDE4A sub-family. A. Co-immunoprecipitations were done using HEK293 cell lysate transiently transfected with POPDC1-myc and PDE4A5-VSV, PDE4B1-VSV or PDE4D7-VSV. Myc-agarose resin was used to precipitate POPDC1 and any interacting PDE4 was detected using VSV tag. Western blotting c for both myc confirmed pull down of POPDC and VSV confirming successful transfection of PDE4 isoforms. Negative control experiments were performed using cell lysate from mock transfected HEK293. B. Specificity of the POPDC1 interaction for endogenous PDE4 isoforms was examined in NRVMs using PanPDE4A, PDE4B and PDE4D antibodies by PLA analysis. C. Images were quantified using ImageJ. Data represents an n of 3 with each n consisting of 15-20 cells. Significance was evaluated using a one-way ANOVA with Tukeys post-hoc analysis, ****p=0.0001, compared to PDE4A control.

### Functional relevance of the POPDC1-PDE4 complex determined by targeted disruption

Since the localization of individual PDE4 isoforms influences their function, the targeted disruption of PDE4 protein-protein interactions can be a highly specific way to investigate the roles of these enzymes (Fertig & Baillie, 2018). Each PDE4 isoform may possess many interaction partners and hydrolyze cAMP at a variety of microdomains within a cell. We hypothesized that a cell-penetrating disruptor peptide that could selectively displace the small fraction of the total PDE4A pool that forms a complex with POPDC1 should have a measurable effect on the function of POPDC1. As we have done for other PDE complexes (Brown *et al*, 2013; Sin *et al*, 2011), we utilized the peptide array information to inform our design of a suitable POPDC1-PDE4A disruptor (Fig. 3) with the sequence TLKKGQTYAAEDKTSVDDRLSILLK (T154-K178 of POPDC1) conjugated to a stearate group to direct trans-membrane transport. A cell-permeable, scrambled peptide was also created using the same amino residues present in the disruptor peptide in a randomized order, and the sequence checked via BLAST to ensure it did not match natural sequences found in other proteins (TTLYTDSSVLKGKRLQDKEKALADI). Co-immunoprecipitations performed with transiently transfected HEK293 cells treated with the disruptor peptide showed a marked reduction in the amount of PDE4A interacting with POPDC1 when compared with those treated with the scrambled control (Fig. 6A). This data suggests that the disruptor peptide was successful in competing against the endogenous PDE4-POPDC1 interaction. In both transiently transfected HEK293 cells (Fig. 6B, C) and NRVMs (Fig. 6D, E) where disruptor peptide treatment (but not scrambled control peptide) significantly reduced the association of PDE4A and POPDC1 detected by PLA.

**Figure 4.**
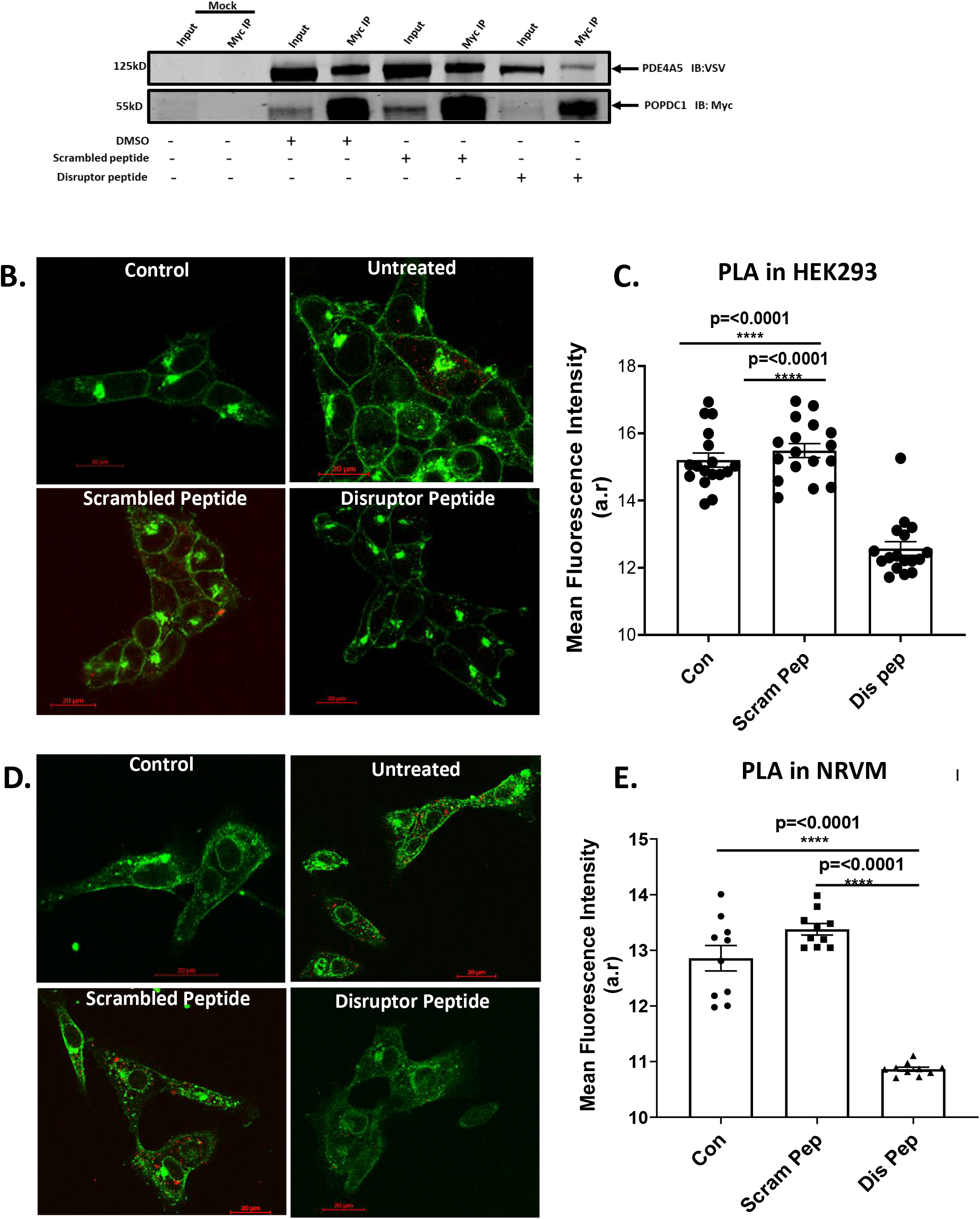
A cell penetrating disruptor peptide dissociates the POPDC1-PDE4 complex. A. HEK293 cells transiently transfected with POPDC1-myc and PDE4A5-VSV were treated with 10 μM scrambled or disruptor peptide. Co-immunoprecipitations of POPDC1 and PDE4A5 performed using Myc-conjugated resin was used to evaluate ability of scrambled or disruptor peptide to compete with PDE4-POPDC1 interaction. The figure is a representative example chosen from three replicate experiments. B. HEK293 cells transiently transfected with POPDC1-myc and PDE4A5-VSV were treated with 10μM scrambled peptide or disruptor peptide prior to fixation. C. After PLA staining of HEK293 cells, images were analyzed using ImageJ-. Significance was evaluated using a one-way ANOVA with Tukeys post-hoc analysis, ****p=0.0001, compared to PDE4A control. D. The experiment was repeated in NRVM staining endogenous proteins. E. After PLA staining of NRVMs, images were analyzed in ImageJ for quantification of PLA signal per condition.. Significance was evaluated using a one-way ANOVA with Tukeys post-hoc analysis, ****p=0.0001, compared to PDE4A control.

As POPDC1 is a cAMP binding protein that binds to and regulates the activity of TREK channels (Schindler *et al.*, 2016), and PDE activity can be modulated by protein and lipid binding partners (Baillie *et al.*, 2019), we set out to determine whether POPDC proteins could regulate the activity of PDE4 enzymes. As expected, the activity of purified PDE4A4 was susceptible to the PDE4 inhibitor rolipram (Fig. Suppl. 1A) but the activity did not change when incubated with different concentrations of purified POPDC1 protein (Fig. Suppl. 1B). POPDC1 did not affect the activity of PDE4A nor did it significantly change the rolipram dose response curve (Fig. Suppl. 2C) suggesting that POPDC1’s association with PDE4 did not occlude the enzyme’s active site. In addition, POPDC1 did not affect the ability of PKA to phosphorylate PDE4A4 as measured with either an antibody to PKA phospho-substrates (Fig. Suppl. 1D,E & F) or against Phospho-UCR1(Fig. Suppl. 1D,E & G). Phosphorylation by PKA activates PDE4 long forms, a mechanism that is partially responsible for the transient nature of cAMP responses (MacKenzie *et al*, 2002). The fact that POPDC1’s interaction with PDE4A does not seem to alter its ability to hydrolyze cAMP suggests that the POPDC1-PDE4A complex is constitutively formed to protect POPDC1 from inappropriate activation under basal conditions. To test this hypothesis we used an established bi-molecular FRET sensor consisting of POPDC1-CFP and YFP-TREK1(Froese *et al.*, 2012) to evaluate the influence of PDE4 activity on the POPDC1-TREK interaction. As expected, increases in global cAMP induced by Forskolin decreased the interaction between POPDC1 and TREK (Fig. 7 A, B). Interestingly, inhibition of PDE4 using the PDE4 specific inhibitor rolipram decreased POPDC1-TREK binding to a greater extent than Forskolin (Fig. 7A, B), suggesting that the PDE4 “pool” that is sequestered to POPDC1 has a highly influential role in “gating” the access of cAMP to PODC1 in order to modulate the POPDC1-TREK interaction.

**Figure 5.**
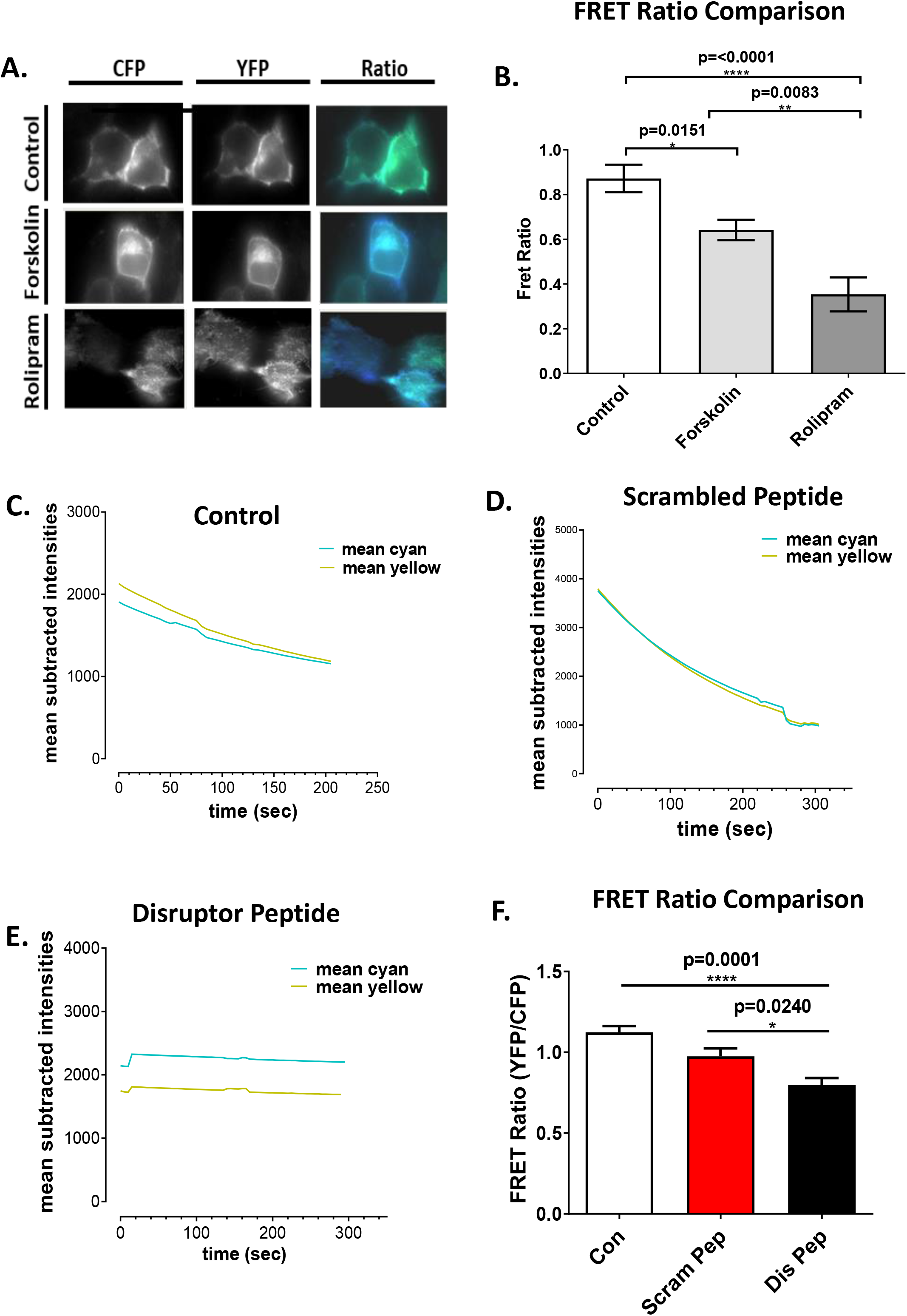
Disruption of the POPDC1-PDE4 complex reduces TREK1 binding. A. HEK293 cells stably expressing PDE4A4 were transiently transfected with POPDC-CFP and TREK1-YFP. After treatment with 25μM Forskolin or 10μM Rolipram, static measurements were taken every 5 seconds for 300 seconds per cell. B. Fret ratio calulated as mean ± SEM, control n=9, forskolin n=15, rolipram n=15 per experiment (n=3). Significance was evaluated using a one-way ANOVA with Tukeys post-hoc analysis, ** p=<0.005, ***p=0.001, compared to control. C. Calculation of the basal FRET ratio (vehicle only treatment) D. Calculation of FRET ratio after treatment with 10μM scrambled peptide or E. 10μM disruptor peptide. F. Static measurements were taken every 5 seconds for 300 seconds. Background fluorescence was subtracted from mean intensities and are plotted as mean subtracted intensities. Results represented as mean ± SEM, untreated n=11, scrambled peptide n=11, disruptor peptide n=20 per experiment (n=3). A one-way ANOVA with Tukey’s post-hoc was used to analyze the FRET ratios.

Next, using the cell-permeable disruptor peptide described above (Fig. 6), we demonstrated that selective disruption of the POPDC1-PDE4 complex could significantly decrease the POPDC1-TREK interaction (Fig. 7 E, F), presumably due to an increase in cAMP binding to POPDC1 resulting from a higher local cAMP concentration in the vicinity of POPDC1 due to the displacement of PDE4A. Treatment with the scrambled peptide control did not produce any significant changes in the POPDC1-TREK interaction as measured by the bi-molecular FRET reporter construct (Fig. 7 C, D, F).

Since the POPDC1-PDE4A disruptor peptide disassociates the POPDC1-PDE4A complex and subsequently interferes with POPDC1-TREK1 interaction, we investigated whether this caused any changes in functional output of ARVMs utilizing the CellOptiq® platform (Clyde Biosciences Ltd; Glasgow, UK). The action potential duration of ARVMs was determined following treatment with the POPDC1-PDE4A disruptor and scrambled control peptides. Measurements of the action potential duration during the action potential repolarization phase (APD 30, 50, 75, 90) were collected from cells which were paced at 1 Hz under baseline conditions (Fig. 7A). No changes were observed at any time point during repolarization for the scrambled peptide in comparison to the untreated control group (Fig. 7B,C,D,E). However, there was a significant elongation of the repolarization phase at APD_30_, APD_50_, APD_75_ and APD_90_ in the presence of the disruptor peptide (one-way ANOVA, post-hoc: Tukey’s multiple comparison p<0.0001) (Fig. 7 B,C,D,E). The prolongation of the APD at 30, 50, 70 and 90 percent suggests that disruption of the POPDC1-PDE4A interaction causes alterations in additional repolarizing currents.

Frequently, changes in action potential duration are met with changes in contractility (Pinnell *et al*, 2007). As such, we sought to determine whether the changes in action potential duration correlated with any alterations in cardiomyocyte contractility. Using the parameter contraction duration 50 (CD_50_), the time between 50% upstroke and 50% downstroke is determined, meaning that both contraction and relaxation are considered. Similar to the APD measurements, there was no alteration between the CD_50_ of myocytes treated with the scrambled peptide cells in comparison to the untreated control cells (Fig. 7F). However, treatment with the disruptor peptide also resulted in no significant changes in CD_50_ time compared to the two control conditions (Fig. 7F). So, although action potentials are elongated, this did not manifest itself as a change in myocyte contractility.

## Discussion

PDE4 activity regulates many crucial cAMP-driven processes in a variety of different tissue types even though their expression level is often low. This apparent contradiction can be explained by the discrete cellular localization of individual PDE4 isoforms, which are sequestered to specific micro-domains in cells by virtue of “postcode” motifs that confer protein-lipid interactions with membranes or protein-protein interactions with a variety of scaffolds. PDE4 containing complexes can be spatially defined or have the ability to translocate to areas of high cAMP levels (Fertig & Baillie, 2018). Once in the correct subcellular domain, PDE4s shape cAMP gradients to allow receptor-specific signals to be relayed via cAMP effector proteins that often exist in close proximity to the phosphodiesterase (Maurice *et al.*, 2014). Aberrant cAMP signaling caused by mutations in cAMP signaling intermediates, PDE4s, their scaffolds or the associated cAMP effector can result in pathologies such as acrodysostosis, ischemic stroke or McCune-Albright syndrome (Jorgensen *et al*, 2015; Kaname *et al*, 2014; Weidner *et al*, 2020).

Here we report for the first time that the most recently discovered cAMP effector, POPDC1, can also be found in a complex with PDE4. Interestingly the binding site for PDE4 on POPDC1 (R172-K178) contains the residue arginine 172 which is mutated to histidine in a family presenting with dilative cardiomyopathy (DCM) (Schindler, 2014). The conclusions from studies investigating the mechanistic effects of this mutation were that cAMP binding affinity of POPDC1 was not significantly affected, nor were there changes in the subcellular localization of the protein or in POPDC1’s ability to interact with TREK1 (Schindler, 2014). Our data suggests the phenotype of the POPDC1^R172H^ mutation may instead be a result of the inability of POPDC1 to bind to PDE4 and thus protect itself from inappropriate cAMP binding under basal conditions. Indeed, in experiments where we tried to phenocopy this situation by using a cell permeable peptide that dislocated PDE4 from POPDC1, we were able to record decreased interaction between POPDC1 and TREK1 (Figure 7) and a prolongation of action potential (Figure 8). Our data provides a putative mechanism that underpins the disease pathology of the POPDC1^R172H^ mutation. Although it is currently not known whether POPDC2 and POPDC3 also bind PDE4, mutation of leucine 155 to histidine in POPDC3, which corresponds to leucine 173 in POPDC1, was recently identified in a patient suffering from limb-girdle muscular dystrophy (LGMDR26, OMIM 605824)(Vissing *et al.*, 2019) Thus, in at least two of the POPDC proteins, mutations in the putative PDE4 interaction motif are associated with heart or skeletal muscle phenotypes.

**Figure 6.**
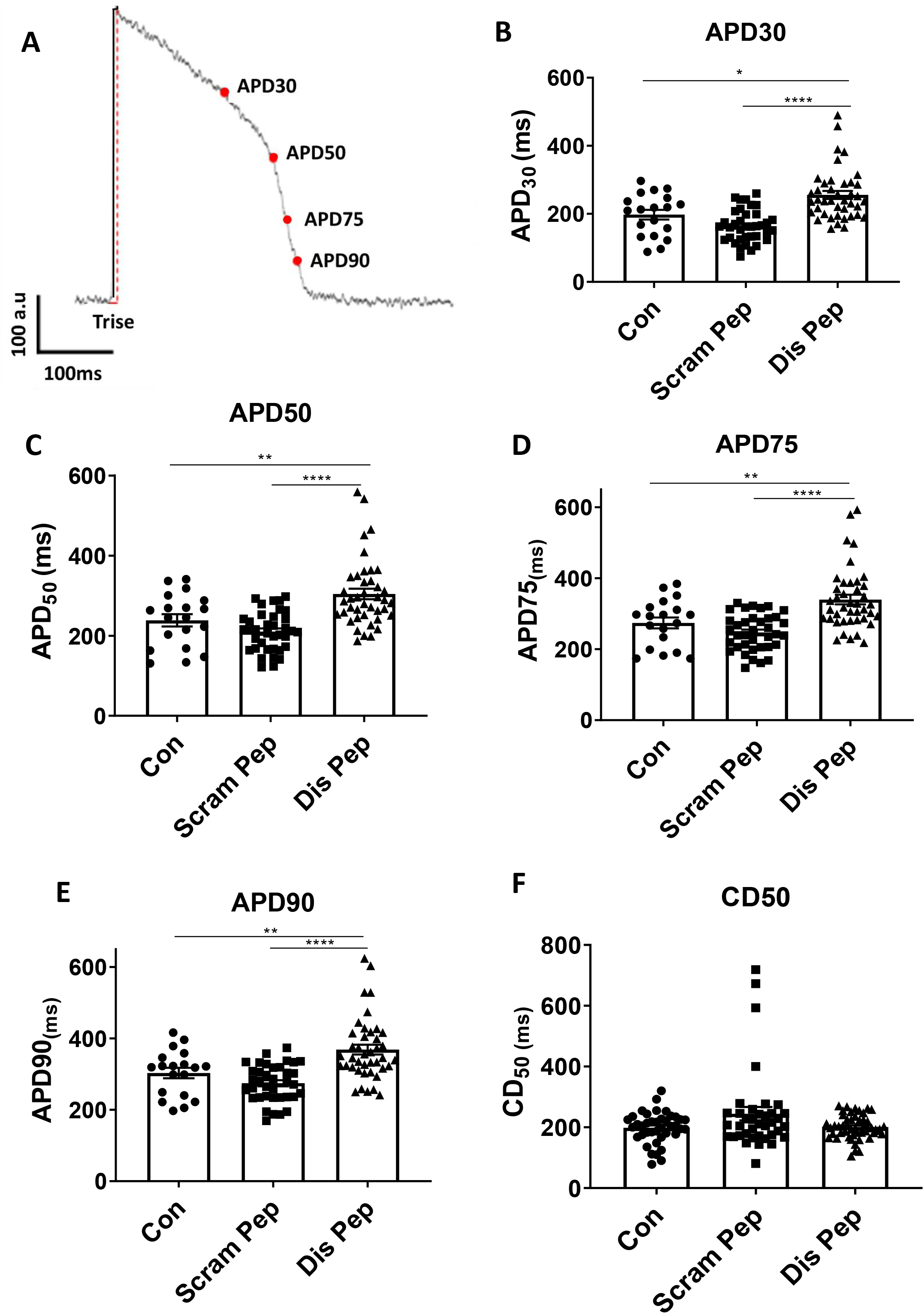
The POPDC1-PDE4 complex influences action potential duration. A. Measurements were collected from 4 points during the repolarization phase. The action potential duration of ARVMs was determined following treatment with the POPDC1-PDE4A disruptor and scrambled control peptides. Measurements of action potential duration across the action potential repolarization phase (B) APD30, (C) APD 50, (D) APD75, and (E) APD90 were collected from cells which were paced at 1 Hz under baseline conditions and displayed on scatter plot graphs with bars ± SEM. Untreated cells, n=19, scrambled peptide n=37 and disruptor peptide n=42 cells. One-way ANOVA with Tukey’s post-hoc analysis, performed using GraphPad Prism™ ****p<0.0001, **p<0.01. E. (F) Contraction duration at 50% contraction to 50% relaxation was measured after 2 hours of treatment with either 10μM scrambled or disruptor peptide. Each point represents the mean value from one cell. Untreated cells, n=19, scrambled peptide n=37 and disruptor peptide n=42 cells. One-way ANOVA with Tukey’s post-hoc analysis, performed using GraphPad Prism™.

**Figure 7.**
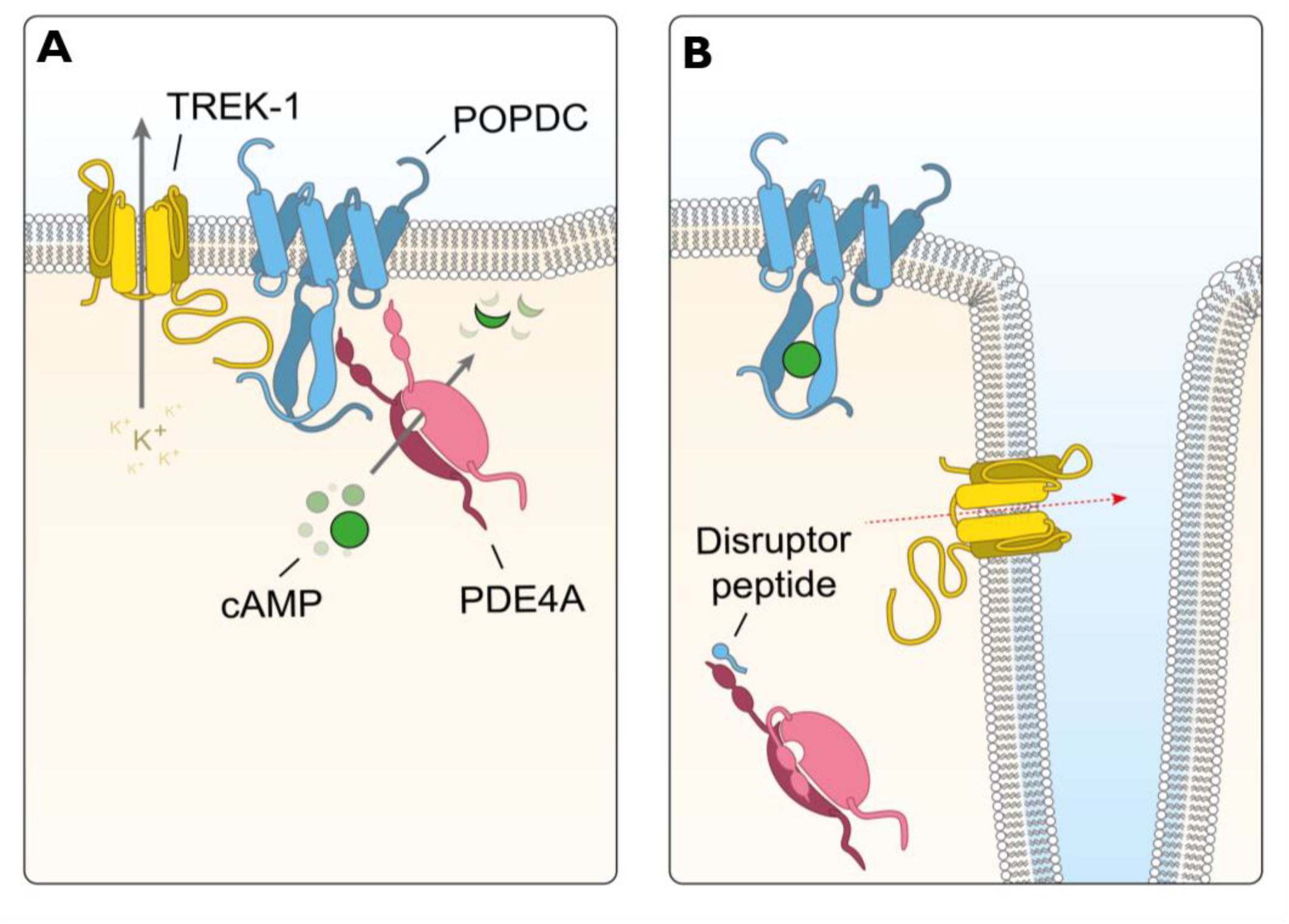
Conceptual summary. **A.** POPDC1 associates with the potassium channel, TREK1 to increase the efflux of K+ ions. This relationship is maintained by local PDE4 activity that keeps cAMP levels low. B. Following dissociation of the POPDC1-PDE4 complex, cAMP readily binds POPDC1 breaking the link with TREK1 to reduce K+ efflux.

Another noteworthy conclusion from our research is that POPDC1 can bind several PDE4 enzymes but has a preference for PDE4As. As the POPDC1 binding site has been mapped to the conserved UCR1 region of PDE4A, it is likely that only the UCR1 containing PDE4 long forms associate with POPDC1. The activity of PDE4 in the human myocardium can be attributed equally to PDE4A and PDE4D isoforms (Johnson *et al.*, 2012) (Richter *et al.*, 2011) and preference for PDE4A binding by POPDC1 must stem from either the ability of both proteins POPDC1 and PDE4A to bind membranes which may bring them into close proximity (Figure 1) or a preferential binding motif within the UCR1 domain of PDE4A compared to other sub-families of PDE4 (PDE4B,C and D (Figure 4). Fine mapping of the POPDC binding site in PDE4 UCR1 identifies a region S162 to S169 (human PDE4A4 numbering) that is completely conserved in all longform PDE4s apart from residues V167 and T168, which are PDE4A specific. This core region must underpin the association with longform PDE4s. The VT di-peptide cassette is only one of the 6 small, divergent, one or two amino acid stretches that are found in that part of UCR1 (Fig.4B). Preliminary data comparing POPDC1 binding to PDE4A vs PDE4D sequences where each highlighted divergent region is sequentially alanised (or Ala to Gly substituted if the amino acid of interest is alanine) shows that PDE4A binds POPDC1 better than PDE4D and is less reliant on the divergent sequences than PDE4D. In fact, when considering each of the six highlighted divergencies between PDE4A and PDE4D, only mutation of the VT dipeptide (and none of the other 5) completely ablates PDE4A binding. This suggests that the preference for PDE4A may lie in this motif. A similar situation exists for the arrestin-PDE4 complex where all PDE4s can associate with the scaffold protein at a site within its N-domain but preference for PDE4D5 is attributed to an additional, specific docking domain in the arrestin C-domain (Baillie *et al*, 2007). Whether PDE4 binding by POPDC1 is shared by the other members of the POPDC family remains an open question. The core sequence identified in POPDC1 is also present in POPDC2 and POPDC3 with some conservative substitutions. It will be interesting to test whether the POPDC2 and POPDC3 also preferentially bind PDE4A, or whether they differ in their binding preference.

The observation that POPDC1 and PDE4 co-localize in heart tissue leaves the possibility that this signaling complex also underpins cAMP transduction events in other tissues open. POPDC1 is expressed most abundantly in striated muscle, i.e. heart and skeletal muscle, and these tissues are affected in patients harboring POPDC1 mutations (De Ridder *et al.*, 2019; Indrawati *et al.*, 2020; Schindler *et al.*, 2016). However, POPDC1 is also found in smooth muscle cells, the gastrointestinal tract, uterus and bladder, neurons of the central and autonomous nervous system and epithelial cells (Andree *et al.*, 2000; Smith & Bader, 2006; Vasavada *et al*, 2004). As PDE4A is also ubiquitously expressed, it is likely that the POPDC1-PDE4 interaction may also be important in other organs and cell types.

The fact that the PDE4-POPDC1 disruptor peptide was able to prolong the repolarization phase of the action potential in ARVM but did not affect contraction duration is surprising as they are usually coupled. However, in agreement with this observation, *Popdc1* null mutant mice display deficiencies in cardiac pacemaking but no effect on cardiac contractility has been reported (Froese *et al.*, 2012). A well-characterized interaction partner of POPDC1 is TREK-1. In co-injected *Xenopus* oocytes, POPDC1 can enhance TREK-1 current, which is thought to be mainly mediated by increased membrane trafficking of the potassium channel (Froese *et al.*, 2012), although effects on channel gating have also been observed (Schindler *et al.*, 2016). The prolongation of the action potential duration after disruptor peptide treatment could at least in part be explained by the modulatory role of POPDC1 on TREK-1(Bodnar *et al*, 2015). Binding of TREK-1 to POPDC1 has been mapped to the proximal part of the Popeye domain (residues 132-186) (Schindler *et al.*, 2016) and thus, is either in proximity or overlap with the PDE4 binding site (residues 172-179). TREK-1 current is potently inhibited by intracellular increases in cAMP via PKA(Patel *et al*, 1998; Terrenoire *et al*, 2001). Two PKA consensus motifs are present in the C-terminus of TREK1, which also binds AKAP79/150 (Sandoz *et al*, 2006). Thus, POPDC1 forming a complex with TREK-1 and PDE4 may either be adjacent to, or part of the AKAP79/150 -PKA complex. POPDC1 could be involved in fine-tuning the effect of adrenergic receptor stimulation on TREK-1 current.

POPDC proteins are multicompartment proteins in cardiac myocytes and are found in cardiac myocytes at the lateral sarcolemma, in t-tubules, caveolae and the intercalated disk (Brand & Schindler, 2017). Moreover, localization at the nuclear envelope and in the nucleoplasm have also been described. It is possible that at each of these subcellular locations different PDE4 isoforms are co-localized with each POPDC, thus modulating the interaction with target proteins.

## Materials and Methods

### HEK-293 cell culture

Human embryonic kidney (HEK) 293 cells were purchased from ATCC and HEK-293 PDE4A4 stable cell line established by Millipore. HEK-293 and HEK-293 PDE4A4 cells were cultured in high-glucose Dulbecco’s Modified Eagle’s Medium (DMEM) supplemented with 10% (v/v) foetal bovine serum, 1% (v/v)penicillin/streptomycin (P/S), 1% (v/v) L-Glutamine, 1% (v/v) minimum essential media non-essential amino acids (NEAA). HEK-293 PDE4A4 cell media is further supplemented with 500 μg/ml G418 (Promega). Culture conditions were 37°C in a humidified atmosphere with 5% CO2. Cells were split at 80-90% confluence and media was replaced every two-three days as required.

### Preparation of Ventricular myocytes from Neonatal Rat and Adult Rabbit

The preparation of neonatal rat cardiomycoytes was modified from the protocol described by in (Bogoyevitch *et al*, 1995). 1-3-day old neonatal Sprague-Dawley rats were sacrificed by cervical dislocation and femoral artery dissection. Cardiomyocytes were dissociated from the ventricles by serial digestions with 0.3 mg/ml Type-2 Collagenase (Worthington) and 0.6 mg/ml pancreatin (Sigma) in a balanced salt solution (120 mM NaCl, 20 mM HEPES, 5.5 mM Glucose, 5.4 mM KCl, 1 mM NaH2PO4 and 0.8 mM MgSO4 (pH 7.4)). Tissue was then incubated for 20 minutes at 37°C in a water bath shaking at 200 cycles per minute. This digestion step was repeated 3-5 times or until the tissue was completely digested. Cell suspensions were collected and pelleted by centrifuging for 5 minutes at 1250 rpm. The pellet was resuspended in 2ml of New-born Calf Serum (NCS). Cells were kept at 37°C in a humidified incubator with an atmosphere of 95% air and 5% CO2. After final digestion the cell suspensions were pooled and centrifuged for. The pellet was resuspended in M1 plating media (67ml D-MEM; 25mM HEPES (Invitrogen), 17.5ml M-199 (Invitrogen), 10ml Horse serum, 5 ml New-born calf serum, 1ml Glutamine 200nM and 0.1ml Penicillin-streptomycin). As non-cardiomyocytes become attached easily, the cells were pre-plated on 10cm2 dishes (Corning) for 2 hours to allow differential attachment of non-myocardial cells. Non-adhered cells were collected and centrifuged at 1250rpm for 5 minutes. Cells were counted and plated at a density of 1 x 106 cells per well of a 6 well plate that was pre-coated with sterile 1% (w/v) gelatin (Sigma-Aldrich). For immunocytochemistry, cells were seeded at 1.5 x 105 on mouse lamin (BD Bioscience) (100 μg/ml per coverslip).

Adult rabbit ventricular myocytes (ARVM) were isolated as described in (Donahue et al., 1998).

### Transfection, Western blotting and Immunoprecipitation

DNA plasmid constructs used for transfection included myc-POPDC1, VSV-PDE4A5, VSV-PDE4A4, VSV-PDE4D7, VSV-PDE4B1, CFP-POPDC1 and YFP-TREK1. Plasmid DNA was transiently transfected into HEK-293 and HEK-293 PDE4A4 Stables using Lipofectamine LTX Reagent (Thermo Fisher Scientific) using manufacturers protocol. Cells were transiently transfected with myc-POPDC1 and one of the PDE constructs or CFP-POPDC1 and YFP-TREK1.

Cellular lysates were prepared in either 3T3 lysis buffer (50 mM NaCl, 50 mM NaF, 25 mM HEPES, 5 mM EDTA, 30 mM sodium pyrophosphate, 10% (v/v) glycerol, 1% (v/v) Triton X-100; pH7.5) or a CO-IP buffer (50 mM Tris; pH8.0, 150 mM NaCl, 2 mM EDTA, 1% (v/v) Triton X-100, 0.25% (w/v) bovine gelatine (Sigma-Aldrich)). Both lysis buffers were supplemented with phosphatase and Complete™ EDTA-free protease cocktail inhibitor tablet (Roche). Protein concentrations of the lysates was determined using the Bradford assay and all samples were equalised for protein concentration.

Proteins were separated using SDS-PAGE (4-12% Bis-Tris gels) and then transferred onto nitrocellulose membranes for Western blotting. For immunoprecipitation, POPDC1(BVES), Myc or VSV were used to immunoprecipitate endogenous, and overexpressed POPDC1 or PDE4 isoform respectively. The resulting immunocomplexes were captured using Protein G sepharose beads (Invitrogen) at 4oC overnight on a rotator. The immunocomplexes were collected by centrifuging at 1000 x g for 5 minutes and washed 5 times in CO-IP lysis buffer. The bound protein complexes were eluted in 2x laemmli buffer and subjected to SDS-PAGE and immunoblotting. Negative controls using isotype-matched IgG from the same species as the antibodies used were performed in order to screen for non-specific binding. Proteins were visualised using appropriated Alexa Fluor Secondary antibodies scanned using the Odyssey Infrared Imaging System (LI-COR Biosciences, UK) for fluorescence detection of the secondary antibodies. Fluorescence signal intensity is quantified using the Odyssey application software (LI-COR Biosciences, UK).

### In vitro pull-down assay

Equal molar concentrations of purified recombinant GST-POPDC1 or His-POPDC1 and MBP-PDE4A4 were mixed in 3T3 lysis buffer [25 mM Hepes, 2.5 mM EDTA, 50 mM NaC1, 50 mM NaF, 30 mM sodium pyrophosphate, 10% (v/v) glycerol, 1% (v/v) Triton X-100, pH 7.5, containing Complete™ EDTA-free protease inhibitor cocktail tablets (Roche)]. Samples were incubated end-on-end with gentle agitation for 1 hour at 4oC. Pre-equilibrated Ni-NTA Superflow resin (Qiagen) or glutathione sepharose resin (Amersham Biosciences) were added to the protein mixture and incubated with gentle agitation for another 1 hour at 4oC. Beads were then sedimented by centrifugation at 10,000 x g for 5 minutes followed by three washes in 3T3 lysis buffer. Proteins were then resolved by SDS-PAGE and immunoblotted using GST (Sigma Aldrich), 6His (Sigma Aldrich) or MBP (Abcam) antibodies.

### SPOT synthesis of peptides and overlay experiments

This was carried as described by us in detail elsewhere (Bolger *et al.*, 2006).

### Immunostaining

For immunofluorescent labelling, cardiac myocytes were plated onto laminin-coated sterile glass coverslips and HEK293 cells were seeded onto laminin-free cover slips. HEK293 cells were transiently transfected with Myc-POPDC1 and VSV tagged PDE4A5, PDE4B1 or PDE4D7. After 24-hour incubation on coverslips the cells were fixed using 4% (w/v) Paraformaldehyde for 1 hour at room temperature with gentle agitation. Coverslips were incubated with Wheat-germ agglutinin for 10 minutes at 37oC to stain the membranes of the cells. Coverslips were washed three times with PBS and blocked for 1 hour at room temperature in blocking buffer (PBS supplemented with 0.5% BSA and 0.25% Triton X-100). Primary antibodies; BVES (Santa Cruz), PDE4A (in-house) for cardiomyocytes and Myc (Abcam) and VSV (Sigma Aldrich) for HEK293, were added at a 1:500 or 1:1000 dilution in blocking buffer to the coverslips and incubated overnight at 4oC in a humidity chamber. After washing three times in PBS, Alexa-Fluor antibodies (Donkey anti-Mouse 488, Goat anti-rabbit 488 and Donkey anti-rabbit 546 (Invitrogen) were added at a 1:200 dilution in blocking buffer. Cells were washed a further three times before being mounted to glass slides using mounting media containing DAPI nuclear stain. Imaging was performed using a Zeiss Pascal laser-scanning confocal microscope (LSM) 510 Meta. Images were acquired with Zeiss LSM Image Examiner and analysed on ImageJ.

### Proximity Ligation Assay

HEK293 cell were transiently transfected with Myc-POPDC1 and one VSV tagged PDE4 isoform; PDE4A5, PDE4B1 or PDE4D7. Neonatal rat ventricular myocytes and adult rabbit myocytes were stained using BVES (Santa Cruz) and an in-house made PanPDE4A, PDE4B or PDE4D antibody. Prior to carrying out the protocol cells were treated with either 10μM disruptor peptide or 10μM scrambled peptide for two hours. To visualise protein-protein interactions in both endogenously and overexpressing cells in situ, Duolink® proximity ligation assay (PLA) was employed using the manufactures protocol.

### Fluorescence Resonance Energy Transfer

CFP-POPDC1 and TREK1-YFP sensors used within this work were produced and trialled by Froese and colleagues (Froese et al., 2012).

HEK293 cells were plated on sterilised 24 mm coverslips (VWR) at a low density to allow for single cells to be measured and clear background regions were available to record. Transient transfections with FRET sensors were carried out 48hour prior to imaging according to previous protocols (Froese et al., 2012).

The coverslip was carefully removed from the culturing dish with watchmaker’s forceps and inserted into a metal ring and sealed securely. Coverslips are washed three times with FRET saline (125 mM NaCl, 5 mM KCl, 1 mM Na3PO4, 1 mM MgSO4, 20 mM HEPES, 5.5 mM glucose, 1 mM CaCl2, pH7.4). A 300 μL bath of FRET saline was applied preventing the cells from drying out. The cells were then visualised on an Olympus IX71 Inverted Microscope under 40x or 60x immersion lenses (Zeiss). Image acquisitions were initiated, and real time measurements were taken. Stimulation with the AC activator, 25 μM forskolin or inhibition by 10 μM Rolipram (PDE4 inhibitor) was carried out by diluting the drug in 300 μL FRET saline to allow for total dispersion. For later experiments, cells were treated with; 10 mM scrambled peptide, 10 mM disruptor peptide or DMSO, 2 hours prior to imaging.

Static measurements were taken in 5 second intervals for a total of 300 seconds to reduce the effects of photobleaching. A beam splitter was used to separate CFP and YFP so that images could be used to obtain a ratio of CFP to YFP in defined areas which had been drawn around single cells as well as clear background. The images were converted to mean intensity values allowing for ratios to be calculated. These ratios were used to determine the change in interaction between POPDC1 and TREK1 in various conditions.

### Cell Optiq

#### Preparation of ARVM for Cell Optiq

Left ventricular cardiomyocytes were isolated from 12-week of New Zealad White Rabbits and plates on glass bottom microwell dishes (35mm petri dish, 14mm Microwell; MatTek®). Cells were allowed to settle for 2 hours prior to each experiment. Cells were treated with 10μM disruptor peptide or 10μM scrambled peptide for 2 hours at 37°C prior to measurements being carried out.

#### Measurements of action potential and contractility

CellOPTIQ® (Clyde Bioscience Ltd; Glasgow, UK) software was used in the collection of high-speed images of contracting single ARVM. For this set of experiments, contractility and action potential parameters were evaluated. Cells were paced with electrodes at 40mV with 20ms duration at a frequency of 1Hz. For each cell a 30 second recording was taken recording at 100 frames per second using a 60x objective lens.

Trace analysis was carried out using CellOPTIQ software. The unfiltered trace was baseline-subtracted, filtered using 3-point, 15 pass filter. The upstroke and repolarisation phases were filtered adaptively to target smoothness and measured by number of curve inflections. Values used in analysis were Trise (upstroke 10%-90% depolarisation) and action potential duration (APD) 30,50,75,90 values, representing time (ms) from upstroke to various degrees of repolarisation. For example, APD50 represents the time from 50% contraction to 50% relaxation. These were plotted to allow visualisation of the repolarisation phase.

Contraction measurements in cardiomyocytes were achieved using measurements of sarcomere length. High-speed (100 frames per second) and high resolution (2048×2048 pixels) video was recorded. Image sampling duration allowed for a minimum of 5 beats to be recorded and subsequently averaged for analysis. Contraction duration (CD50) was recorded, considering both contraction and relaxation. CD50 represents the time from 50% contraction to 50% relaxation of the myocyte.

For measurement analysis, the macro programme SarcomereLength created in ImageJ by Dr Francis Burton (University of Glasgow), was utilised. Each stack of images was analysed by manually positioning a linear selection perpendicular to the cell allowing for the capture of isotropic contractions. The line length was dependent upon cell size and width was kept constant at 50 pixels to reduce any pixel noise.

#### cAMP Phosphodiesterase Assay

Recombinant purified MBP-PDE4A4 was incubated with GST-POPDC1 for 30 minutes prior to the start of this protocol. MBP-PDE4A4 was used consistently at 10μg for all experiments with the concentration of GST-POPDC1 was increased from 0 to 50μg. Samples were treated with increasing concentrations of Rolipram from 0.5μM to 10μM. The cAMP PDE assay was then carried out as described in (Marchmont & Houslay, 1980)

## Acknowledgements

GSB is supported by BHF PG/17/26/32881and AJT by the College of Veterinary, Medical and Life Sciences Doctoral training Programme.

## Author contributions

GSB, BS and AJT conceived the project and wrote manuscript with input from TB, WF and GS. AJT undertook the majority of experimental work with help from AMC, CG, CB, GST and RM. GLS, SD and NM planned and carried out all functional myocyte experiments.

## Author contributions

GSB, BS and AJT conceived the project and wrote manuscript with input from TB, NM and GS. AJT undertook the majority of experimental work with help from AMC, CB, GST and RM.

---

**supplementary 1.**
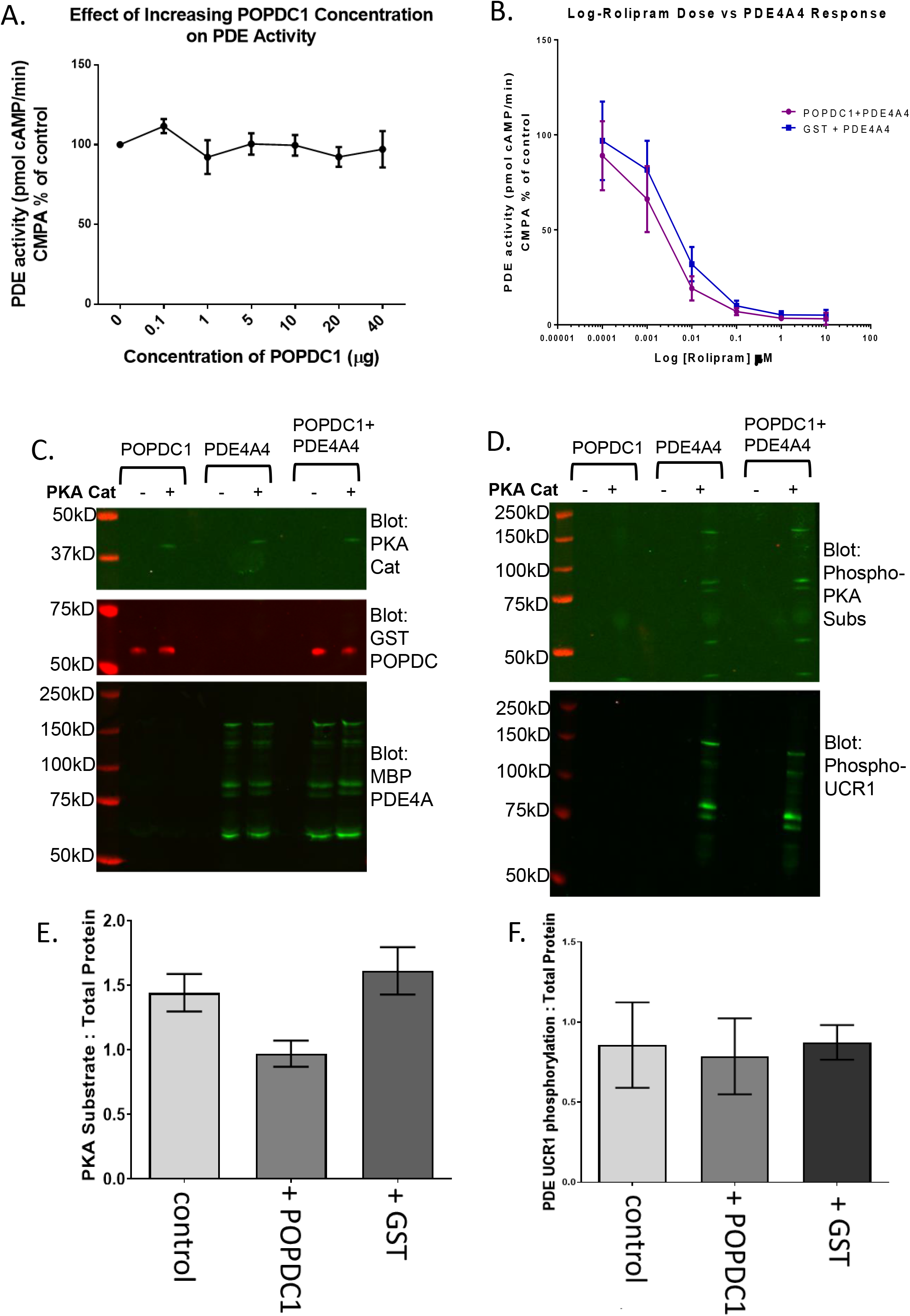

